# A novel conceptual model of heart rate autonomic modulation based on a small-world modular structure and heterogeneous innervation of the sinoatrial node

**DOI:** 10.1101/2023.08.06.552167

**Authors:** Alexander V Maltsev, Michael D Stern, Edward G Lakatta, Victor A Maltsev

**Author notes:** **Corresponding author:** Victor A. Maltsev.

## Abstract

The current theory of cardiac pacemaker rate modulation by the autonomic nervous system is based on the concept that a primary pacemaker cell or a group of cells in the center of the sinoatrial node (SAN) can change its AP firing rate within a broad range, driving respective myocardium contractions to commensurate body demands. Experimental data show, however, that pacemaker cells are extremely heterogeneous, with different areas of the SAN or cell clusters specializing to drive APs at specific rates. Thus, higher heart rates under stress are mainly driven by cell clusters in superior SAN, whereas low rates by cell clusters in inferior SAN, with basal state rates generated somewhere in the middle of the node. Cells within different clusters feature different intrinsic electrophysiological and Ca cycling properties, sympathetic and parasympathetic innervation, and vasculature, thereby supporting effective shift of the system to an optimal rate (under given conditions) accompanied by respective shifts in the leading pacemaker site. Thus, the popular single-cell-based pacemaker theory does not capture this complex emerging paradigm of pacemaker function of SAN tissue revealed by recent experimental studies. Here we propose a more realistic, conceptual model of heart rate autonomic modulation based on these studies. Our new model (the ‘gear model’) simulates the SAN as a brain-like structure featuring a small world of loosely connected clusters (functional modules) of tightly coupled cells, modeled as Maltsev-Lakatta coupled-clock system. One module (the higher chronotropic gear) generates higher AP rates in basal state and under β-adrenergic stimulation, but its activity is strongly suppressed by parasympathetic stimulation. The other module (the lower gear) generates lower rates and has low sensitivity to parasympathetic stimulation. Such modular, gear-like system reproduces the respective shifts of the leading pacemaker site observed experimentally and features a wide range of rate modulation and robust function whilst conserving energy. In perspective, future refinement and application of this new pacemaker tissue mechanism will provide better understanding of cardiac pacemaker function, its deterioration in aging and disease, and ultimately the creation of new therapies to treat sick sinus syndrome and other SAN function-related cardiac arrythmias.

## INTRODUCTION

The sinoatrial node (SAN) is the primary heart pacemaker, generating rhythmic electrical impulses drives heart contractions at rates and strength to satisfy blood supply under given condition. Since the successful application of Hodgkin-Huxley theory to heart cells (Noble, 1960), the cardiac pacemaker field had been long dominated by the idea that the ensemble of membrane ion currents (dubbed later as the membrane clock (Maltsev and Lakatta, 2009)) drives the diastolic depolarization and spontaneous AP firing in pacemaker cells. Primary pacemaker cells in the SAN center were thought to dictate the excitation rate of other SAN cells (Sano et al., 1978;Bleeker et al., 1980). Thus, extensive search for the mechanisms of autonomic modulation of heart rate had initially been mainly limited to respective modulation of specific membrane ion currents, such as *I*_CaL_, *I*_K_, *I*_f_, *I*_KACh_ in single cell models (Hauswirth et al., 1969;Brown et al., 1975;Noma and Trautwein, 1978;Brown et al., 1979;DiFrancesco et al., 1989;Demir et al., 1999;Zhang et al., 2002;Himeno et al., 2008;Tao et al., 2011). Further studies revealed critical importance of Ca cycling in cardiac automaticity, known as Ca clock (Lakatta et al., 2003;Maltsev et al., 2004), the coupled clock system (Maltsev and Lakatta, 2009), and more recently as the ignition process (Lyashkov et al., 2018). Critical importance of local Ca releases for β-adrenergic receptor (βAR) and cholinergic receptor (ChR) stimulations have been demonstrated by live-cell confocal microscopy imaging (Vinogradova et al., 2002;Lyashkov et al., 2009). Thus, the search for numerical formulations for autonomic modulation mechanisms included the coupled-clock function, but still within the single pacemaker cell paradigm (Vinogradova et al., 2006;Lyashkov et al., 2009;Maltsev and Lakatta, 2010; 2013).

The SAN cell population, however, is extremely heterogeneous with respect to cell shape, size, and biophysical properties. The expression of Ca cycling proteins and membrane ion channels vary substantially among individual SAN cells (Honjo et al., 1996;Honjo et al., 1999;Lei et al., 2001;Musa et al., 2002;Monfredi et al., 2017). For example, *I*_*CaL*_ density varies by an order of magnitude (Monfredi et al., 2017) and cells differ dramatically by their response to autonomic modulation (Kim et al., 2021;Yang et al., 2021). Furthermore, some cells isolated from SAN (dubbed dormant cells) do not generate spontaneous AP firing but can “awake” and generate rhythmic APs during βAR stimulation (Kim et al., 2018;Louradour et al., 2022). Cells isolated from superior or inferior regions of SAN also exhibit different automaticity (Grainger et al., 2021). Heterogeneous cell properties are in agreement with recent results of high-resolution imaging of intact SAN tissue at a single cell level. These studies demonstrated that while the majority of SAN cells indeed fire synchronously with a common period, many cells fire at various rates and irregularly, or remain silent, generating only local Ca releases (Bychkov et al., 2020;Fenske et al., 2020), like dormant cells discovered previously in single cell studies. The number of silent cells in tissue increased by ChR stimulation (Fenske et al., 2020). Overall, synchronized AP firing seems to emerge from heterogeneous signalling with the heart’s pacemaker mimicking brain cytoarchitecture and function, thereby resembling multiscale complex processes of impulse generation within clusters of neurons in neuronal networks (Bychkov et al., 2020;Bychkov et al., 2022). Opthof (Opthof, 2007) stated that “normal sinus node remains spontaneously active at high concentrations of acetylcholine, because it has areas that are unresponsive to acetylcholine”. The SAN indeed exhibits heterogeneous autonomic innervation, with some regions having extremely low parasympathetic innervation (e.g. ∼10% in epithelial vs. endothelial regions) (Bychkov et al., 2022).

Application of neurobiochemical marker S100B (Ca-binding protein B) to the SAN desynchronized Ca signalling revealed individual cell clusters operating at lower rates (Bychkov et al., 2022). Furthermore, two spatially distinct competing pacemaker regions, superior and inferior SAN (termed as sSAN and iSAN) have been identified in the mammalian hearts (Brennan et al., 2020). Those regions feature different profiles of expression of ion channels, cardiac receptors, neural proteins, and transcription factors and preferentially control the fast and slow heart rates via autonomic nervous system modulation that accompanied by respective shift of leading pacemaker locations (Brennan et al., 2020). The pacemaker shift caused by autonomic modulation has been reproduced in a variety of numerical models of SAN tissue. However the SAN tissue in those models was populated by heterogeneous cells either randomly (Syunyaev and Aliev, 2011) or following a gradient distribution (Munoz et al., 2011;Inada et al., 2014). On the other hand, the shift is likely “linked to the presence of distinct anatomically and functionally defined intranodal pacemaker clusters that are responsible for the generation of the heart rhythm at different rates” as recently hypothesized by (Lang and Glukhov, 2021) based on substantial experimental data in mouse, rabbit, canine and human SAN.

Thus, recent research indicates that the existing single cell-based models of autonomic modulation of the heart rate are largely inadequate to explain cardiac pacemaker automaticity at the tissue level. At the same time, existing tissue models do not include function of pacemaker cell clusters observed experimentally. While small clusters of cells can emerge spontaneously in randomly distributed cells (Maltsev et al., 2022a) due to Poisson clumping (Aldous, 1989), the specialized pacemaker clusters are actually much larger, e.g. sSAN and iSAN (Brennan et al., 2020), and, as such, likely emerge via morphogenesis rather than randomly.

Here we propose a novel conceptual numerical model of heart rate autonomic modulation based on recent experimental studies in SAN cells in tissues. Our model simulates the heterogeneous SAN as a network of loosely connected clusters (modules) consisting of tightly coupled cells, featuring clustered small world topology, like in brain networks (Russo et al., 2014;Bassett and Bullmore, 2017). Each module, in turn, specializes to drive its specific rate range, like gears in a car. In our simple model, a higher gear module generates higher AP rates in basal state and under β-AR stimulation. Its activity is strongly suppressed by parasympathetic stimulation, while a lower gear module, insensitive to ACh, drives the system at its low rate.

## METHODS

Each pacemaker cell in our multicellular SAN model was featured in the Maltsev-Lakatta model of central SAN cells, having a coupled-clock pacemaker mechanism (Maltsev and Lakatta, 2009). The computer code for the original model can be freely downloaded in CellML format at http://models.cellml.org/workspace/maltsev_2009 and executed using the Cellular Open Resource (COR) at http://www.opencor.ws/. The Supplementary text provides further details, including formulations of βAR and ChR stimulation. Supplementary Table S1 provides the initial conditions. Computations were performed using NVIDIA RTX A6000 GPU as previously described (Maltsev et al., 2022a), based on the original CUDA-C algorithm of (Campana, 2015). All cells were identically connected with ρ = 3750 MΩ*m.

## RESULTS

### Developing new multicellular model

We developed a new computational algorithm in Python that allows for SAN network to evolve from seeding points following a given distribution of connections (https://github.com/alexmaltsev/IterativeNetworkGenerator). Our algorithm iteratively creates a network of loosely connected clusters (modules) based on circular probability mass function attachment preference with two variables for interconnections and intraconnections. While the exact connectome for SAN has not yet been established experimentally, we assume here a reasonable number of connections between neighboring cells ranging from 1 to 5 (Fig. 1A) based on visual inspection of previously reported high-resolution confocal images of SAN tissue (Bychkov et al., 2020;Bychkov et al., 2022). A representative example of the modular system is illustrated in Figure 1B and in Supplementary Figure S1 in computer-mouse-interactive 3D viewer.

**Figure 1.**
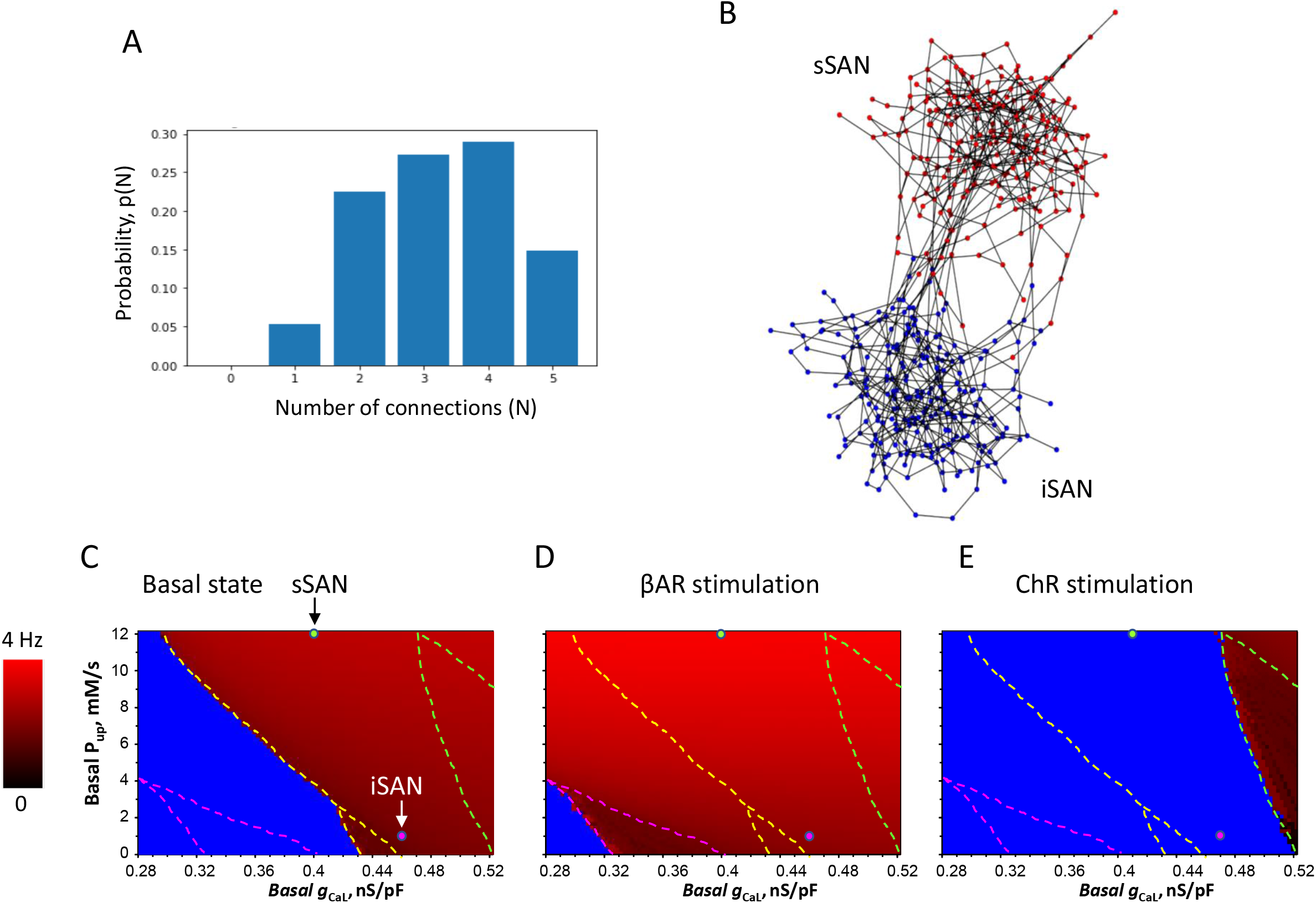
Development of 3D model of SAN with brain-like modular structure and function. A: Distribution of number of connections between cells used for creation of SAN functional modules. B: An example of the model consisting of two modules: 242 sSAN cells (red circles) and 243 iSAN cell (blue circles). A detailed interactive 3D view of the model is provided in Supplementary Figure S1. C-E: results of parametric sensitivity analysis of βAR and ChR stimulation based on the basal conductance of *I*_CaL_ (*g*_CaL_, *x* axis) and SR Ca pumping rate (*P*_up_, *y* axis). The image shows the result of 97 (*x*)*61 (*y*) = 5,917 simulations (each pixel =0.0025 nS/pF * 0.2 mM/s). The fork-like dash lines outline the bifurcation borders between non-firing (blue), firing (gradual red shades) and chaotic firing (mosaic red shades within lower part of the fork). Red shades code AP firing rates from 0 (black) to 4 Hz (pure red). The sSAN and iSAN were populated with cells with parameters marked by green and magenta circles, respectively.

### Parametric sensitivity analyses of cell parameters

For simplicity and clear interpretation, the cell parameters were differentially set among modules, but identical within each module, tuned to generate specific rates: 1) the superior module (sSAN, as termed in (Brennan et al., 2020)) had ion currents and SR Ca pumping rate to generate physiological AP firing rates in basal state and during βAR stimulation; 2) the inferior module (iSAN) had parameters to generate lower physiological AP rates during ChR stimulation.

While we used the same formulations to model βAR stimulation in each module, ChR stimulation was applied only to sSAN. We performed wide-range sensitivity analyses (Fig. 1C-E) with respect to two key model parameters, basal *g*_CaL_ determining maximal *I*_CaL_ conductance, and basal *P*_up_ determining maximal SR Ca pumping rate for three conditions: 1) basal state; 2) βAR stimulation; 3) ChR stimulation. Cell parameters were chosen for sSAN to operate with a strong contribution of Ca clock (*P*_up_ =12 mM/s) and a moderate *g*_CaL_ of 0.4 nS/pF (green circle) to reproduce experimental result of cessation of AP firing in isolated sSAN under ChR stimulation (Brennan et al., 2020) (Fig. 1E). Cell parameters were chosen for iSAN to operate mainly as membrane clock (*P*_up_ =1 mM/s, *I*_CaL_=0.46 nS/pF, magenta circle) in close proximity to the bifurcation border to generate lowest possible AP rate without arrhythmia when the sSAN is suppressed by ChR stimulation.

### Autonomic modulation of the system

In the basal state and during βAR stimulation the leading module was sSAN firing APs with mean rates of 3.0 and 3.9 Hz, respectively, while the iSAN fired APs with a substantial delay and at lower mean rates of 2.2 and 2.6 Hz, respectively (Fig. 2A,B, Supplementary Videos 1 and 2). Next, we simulated the system behavior in ChR stimulation. In this case iSAN (insensitive to ACh) became the leading module generating APs at a low mean rate of 2.08 Hz (Fig. 2C, Supplementary Video 3). While sSAN could not generate APs in isolation in the presence of ACh (green circle in Fig. 1E), it was paced by iSAN within the system (through the loose inter-modular connection) to generate AP at about the same rate, albeit with a substantial delay. Thus, the modular system was able to generate the entire range of physiological AP firing rates (for rabbit) from 2.08 to 3.9 Hz by two loosely coupled modules, each of which was tuned for different rates with appropriate innervation.

**Figure 2.**
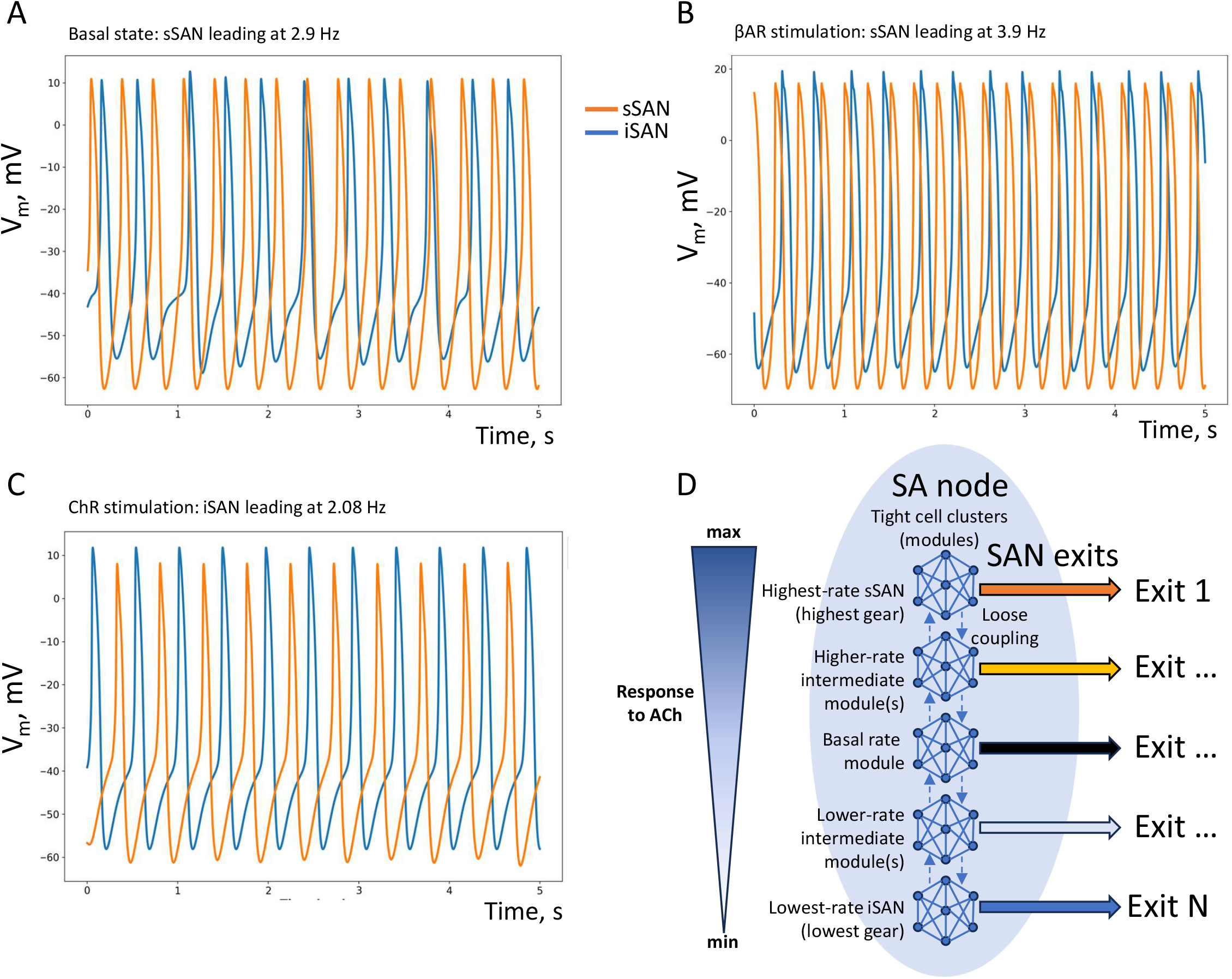
Performance of the “gear” model in different conditions. A, B, C: simulated AP traces in basal state (in the absence of autonomic modulation), in βAR stimulation, and ChR stimulation [ACh], respectively, in each pacemaker module: sSAN (orange traces) and iSAN (blue traces). The respective leading/pacing rates are shown at the top of each panel. D: Schematic diagram of our new conceptual model (the “gear” model) of heart rate autonomic modulation based on brain-like modular structure and heterogeneous innervation of the SAN.

## DISCUSSION

Theories of autonomic modulation of heart rate have evolved thus far at the single cell scale following the idea that the primary pacemaker cell in the center of SAN generates APs over the entire physiological rate range and paces other cells. This approach does not capture the modern view on SAN structure-function relationship gleaned from experimental studies, i.e. distinct anatomically and functionally defined intranodal pacemaker clusters generate the heartbeats at different rates (Brennan et al., 2020;Bychkov et al., 2020;Lang and Glukhov, 2021). Here, for the first time, we provide a numerical validation of the concept that the entire range of autonomic modulation of heart rate can be achieved via brain-like structure featuring loosely connected clusters (functional modules) of tightly coupled cells, specialized for particular AP firing rates.

In such a system, as in the brain (Russo et al., 2014;Bertolero et al., 2015;Zimmern, 2020), the maximum efficiency is achieved when the system operates at the edge of criticality (Fig. 1C-E). Many biological systems benefit from operating at the edge of criticality to reap benefits such as “an optimal balance between robustness against perturbations and flexibility to adapt to changing conditions as well as to confer on them optimal computational capabilities, large dynamical repertoires, unparalleled sensitivity to stimuli, etc.” (cited from (Muñoz, 2018)).

Based on these considerations and our new model simulations, the role of parasympathetic stimulation in the new emerging paradigm of heart rate modulation is not to decrease the AP rate of the pacemaker cell per se, but rather to suppress the activity of module(s) generating higher AP rates in order to unmask the function of other module(s) insensitive (or less sensitive) to ACh which are tuned to safely operate at lower rates. Thus, the SAN pacemaker system is modulated via anatomically and functionally distinct cell clusters (Brennan et al., 2020;Bychkov et al., 2020;Lang and Glukhov, 2021), similar to switching car gears to achieve the best performance at different speeds under different conditions. In these terms, the ChR stimulation acts as a powerful downshifter, overriding βAR stimulation in agreement with well-known phenomena known as anti-adrenergic effect (Belardinelli et al., 1995;Norton et al., 1999), i.e. stronger ACh and adenosine effects in the presence of βAR stimulation, as well as accentuated antagonism, i.e. substantially smaller sympathetic heart rate effects at high levels of vagal tone (Levy, 1984;Mizuno et al., 2008)).

One important benefit of such modular system is energy saving. Indeed, while the membrane clock of iSAN still generates rare APs under parasympathetic stimulation, intrinsic automaticity of sSAN becomes completely suppressed (Brennan et al., 2020), i.e. it does not consume energy for its Ca clock operation which is the major part of energy budget in SAN cells (Yaniv et al., 2011).

With respect to limitations and future studies, we present here proof of principle for the new modular pacemaker mechanism, assuming that numerous missing details of anatomical structures and heterogeneity of cell biophysical properties within each module (Honjo et al., 1996;Honjo et al., 1999;Lei et al., 2001;Musa et al., 2002;Monfredi et al., 2017) will be implemented in future model upgrades. For example, the cell model (Maltsev and Lakatta, 2009) used here cannot generate very low basal AP rates below 2Hz (magenta point in Fig. 1C) but this can be improved if some inward currents (e.g. *I*_st_, *I*_bNa_) with unknown molecular identity are excluded, allowing to reach AP rates as low as 1 Hz (Maltsev and Lakatta, 2013). A major model upgrade could be implementation of heterogeneous, local Ca releases (Stern et al., 2014;Maltsev et al., 2022b). Further, recently reported different profiles of local expression of ion channels, cardiac receptors, neural proteins, and transcription factors in different regions of SAN (Brennan et al., 2020) will be very helpful in further model upgrades.

While our study was inspired by the idea of SAN operating as a neuronal network (Bychkov et al., 2020;Bychkov et al., 2022) and by the discovery of sSAN/iSAN by (Brennan et al., 2020), an important next step will be to quantify SAN connectome and physiome to generate more realistic 3D-structure of the functional modules. For example, modular systems are small-world but not all small-world systems are modular. “Brain networks and many other complex systems demonstrate the property of hierarchical modularity, or modularity on several topological scales: within each module there will be a set of sub-modules, and within each sub-module a set of sub-sub-modules, etc. There are several general advantages to modular and hierarchically modular network organization, including greater robustness, adaptivity, and evolvability of network function” (cited from (Meunier et al., 2010)). While the physical nature of weak functional connections of HCN4+/CX43-cells in SAN center remains unknown, recent computational studies showed that pacemaker function is possible only within a specific range of cell connectivity (not too tight and not too loose); and cell heterogeneity substantially increases robustness of the system operation (Campana et al., 2022;Maltsev et al., 2022a;Maltsev and Stern, 2022).

An obvious future model upgrade could be also adding intermediate chronotropic “gears”, including “basal state” module (Fig. 2D). This further module specialization would likely provide further energy savings and more efficient operation at a given condition.

Furthermore, each SAN has several exits to rhythmically excite atria that is important for robust delivery of SAN impulses to atria (Li et al., 2017;Easterling et al., 2021). Thus, upgraded further models will add cellular link(s) to CX43+ network and the SAN exits. An interesting hypothesis could be then that each such exit is associated with the respective “chronotropic gear” of the module fine-tuned for particular rate range (Fig. 2D). We have shown that formal clustering of cells with similar properties increases robustness of the system to generate APs at the edge of criticality (Maltsev et al., 2022a); but the problem was to exit APs outside the relatively small clusters to stimulate atria. The modular system organization of one-exit-per-module will likely solve this problem. Also, the system of sequential loosely coupled clusters (Fig. 2D) is likely less disposed to reentrant arrhythmias, due to its very structure lacking reentrant pathways among the modules.

An interesting idea of SAN operation via a percolation phase transition was proposed by (Weiss and Qu, 2020). In our model, such percolation would easily occur within each cluster of tightly connected cells but not through all loosely connected clusters, keeping their fine-tuned specific rates intact, thereby delivering robust and flexible operation of the gear system. Moreover, the loose connections link the leading cluster to the neighboring cluster and its neighboring (redundant) exit that can ensure robust SAN operation in case the exit of the leading cluster is blocked.

The SAN is controlled by so called the heart’s little brain (Armour, 2008;Herring and Paterson, 2021). The results of the present study support and further develop this idea by adding theoretical insights into efficient regulation of SAN network operating as brain-like modular structure. The entire SAN system is not limited to SAN pacemaker cell network. In addition to heterogeneous innervation (modelled here), recent studies identified autonomic plexus, peripheral glial cell web, and a novel S100B(+)/GFAP(–) interstitial cell type embedded within the HCN4+ cell meshwork that all increase the structural and functional complexity of the SAN and provide a new regulatory pathway of rhythmogenesis (Bychkov et al., 2022). Thus, additional layers of cellular networks of different nature are likely to be very important to tune and keep the modular system at its optimal performance. For example, the energy benefit of the modular structure (captured by the present model) can be further enhanced with other mechanisms, not considered here: 1) presence of energy-saving dormant cells (Kim et al., 2018;Bychkov et al., 2020;Fenske et al., 2020;Louradour et al., 2022;Maltsev and Stern, 2022) and 2) higher vascular density in sSAN places pacemaker cells with metabolically demanding, high-rate APs near vessels, i.e. new blood supply (Grainger et al., 2021).

As the modular structures are essential to memory (Rodriguez et al., 2019), switching between chronotropic gears may contribute to complex patterns of healthy heart rate variability. Loss of heart rate variability or its complexity is linked to age- and disease-related decline in cardiovascular health. A general tendency in aging is the loss of complexity of biological structures, leading to the age-related decline in adaptive capacity (Lipsitz and Goldberger, 1992). Therefore, possible loss of the complex modular organization of SAN tissue could explain, in part, age-related decline in SAN function (including age-dependent decline in maximum heart rate): the system may lose its distinct gears and efficient energy operation. Such unstructured system would be also prone to emergence of reentrant arrythmia and chaotic firing in the random mixture of cells with different properties (Maltsev et al., 2022a).

## SUPPLEMENTARY MATERIAL

The Supplementary Material includes 3 Supplementary videos, Supplementary text providing mathematical formulations, Supplementary Figure S1 showing our model in interactive 3D viewer, Supplementary Table S1 with initial values of all 29 variables (identical in each cell).

### Legends for Supplementary videos

**Video 1.** Membrane potential dynamics in each cell of the SAN model with loosely coupled modules in basal state. *V*_*m*_ values were color-coded via red shade from -60 mV (pure black) to 20 mV (pure red). Simulation is 5 s long. sSAN leads pacing.

**Video 2.** Membrane potential dynamics in each cell of the SAN model with loosely coupled modules during βAR stimulation. The same system and color coding as in Video 1. sSAN leads pacing.

**Video 3.** Membrane potential dynamics in each cell of the SAN model with loosely coupled modules during ChR stimulation. The same system and color coding as in Video 1. iSAN leads pacing.

