## Supplementary text for "A novel conceptual model of heart rate autonomic modulation based on a small-world modular structure and heterogeneous innervation of the sinoatrial node"

**DESCRIPTION OF SINGLE CELL MODEL AND ITS AUTONOMIC MODULATION**

We performed simulations using modified Maltsev-Lakatta central SAN cell numerical model that features a coupled-clock mechanism (Maltsev and Lakatta, 2009). The original model is freely available and can be downloaded and run in CellML format (<http://models.cellml.org/workspace/maltsev_2009>) using the Cellular Open Resource software developed by Alan Garny at Oxford University in the UK (Garny et al., 2009) (for recent development of this software see <http://www.opencor.ws/>). We simulated APs in three scenarios: basal AP firing, β adrenergic receptor (βAR) stimulation with 0.1 μM of isoproterenol, and cholinergic receptor (ChR) stimulation with acetylcholine (ACh), 0.1 μM.

**MODEL PARAMETERS**

**Fixed ion concentrations, mM**

*Ca*_o_ = 2: Extracellular Ca^2+^ concentration.

*K*_o_ = 5.4: Extracellular K^+^ concentration.

*K*_i_=140: Intracellular K^+^ concentration.

*Na*_o_ = 140: Extracellular Na^+^ concentration.

*Na*_i_=10: Intracellular Na^+^ concentration.

*Mg*_i_ = 2.5: Intracellular Mg^2+^ concentration.

**Cell compartments**

*C*_m_ = 32 pF: Cell electric capacitance.

*L*_cell_ = 70 μm: Cell length.

*R*_cell_ = 4 μm: Cell radius.

*L*_sub_ = 0.02 μm: Distance between jSR and surface membrane (submembrane space).

*V*_cell_ = π·*R*_cell_^2^·*L*_cell_ = 3.5185838 pL: Cell volume.

*V*_sub_ = 2π·*L*_sub_·(*R*_cell_ - *L*_sub_/2)·*L*_cell_ = 0.035097874 pL: Submembrane space volume.

*V*_jSR_part_ = 0.0012: Part of cell volume occupied by junctional SR.

*V*_jSR_ = *V*_jSR_part_·*V*_cell_: Volume of junctional SR (Ca^2+^ release store).

*V*_i_part_ = 0.46: Part of cell volume occupied with myoplasm.

*V*_i_ = *V*_i_part_·*V*_cell_-*V*_sub_: Myoplasmic volume.

*V*_nSR_part_ = 0.0116: Part of cell volume occupied by network SR.

*V*_nSR_ = *V*_nSR_part_·*V*_cell_: Volume of network SR (Ca^2+^ uptake store).

**The Nernst equation and electric potentials, mV**

E_X_ = (RT/F) · ln([X]_o_/[X]_i_) = *E*_T_ · ln([X]_o_/[X]_i_), where

F = 96485 C/M is Faraday constant,

T = 310.15 K˚ is absolute temperature for 37˚C,

R = 8.3144 J/(M·K˚) is the universal gas constant,

*E*_T_ is “RT/F” factor = 26.72655 mV,

and [X]_o_ and [X]_i_ are concentrations of an ion “X” out and inside cell, respectively.

*E*_Na_ = *E*_T_ ∙ ln(Na_o_/Na_i_): Equilibrium potential for Na^+^.

*E*_K_ = *E*_T_ ∙ ln(K_o_/K_i_): Equilibrium potential for K^+^.

*E*_Ks_ = *E*_T_ ∙ ln[(K_o_ + 0.12∙ Na_o_)/(K_i_ + 0.12 ∙ Na_i_)]: Reversal potential of *I*_Ks_.

*E*_CaL_ = 45: Apparent reversal potential of *I*_CaL_.

*E*_CaT_ = 45: Apparent reversal potential of *I*_CaT_.

*E*_st_ = 37.4: Apparent reversal potential of *I*_st_.

**Sarcolemmal ion current types and their parameter values**

*I*_CaL_:	L-type Ca^2+^ current (*g*_CaL_ = 0.464 nS/pF).

Steady-state activation parameters: *V*_½,d_ =-13.5 mV; *K*_d_ =6 mV.

Steady-state inactivation parameters: *V*_½,f_ =-35 mV; *K*_f_ =7.3 mV.

*K*_mfCa_ = 0.00035 mM: Dissociation constant of Ca^2+^ -dependent *I*_CaL_ inactivation.

*β*_fCa_ = 60 mM^-1^ · ms^-1^: Ca^2+^ association rate constant for *I*_CaL_.

*α*_fCa_ = 0.021 ms^-1^ : Ca^2+^ dissociation rate constant for *I*_CaL_

*I*_CaT_:	T-type Ca^2+^ current (*g*_CaT_ = 0.1832 nS/pF).

*I*_f_:	Hyperpolarization-activated current (*g*_If_ = 0.15 nS/pF).

*V*_If,1/2_ = -64 mV: half activation voltage for *I*_f_ current in the basal state.

*I*_st_:	Sustained non-selective current (*g*_st_= 0.003 nS/pF).

*I*_Kr_:	Delayed rectifier K^+^ current rapid component (*g*_Kr_ = 0.08113973 nS/pF).

*I*_Ks_:	Delayed rectifier K^+^ current slow component (*g*_Ks_ = 0.0259 nS/pF).

*I*_to_:	4-aminopyridine sensitive transient K^+^ current (*g*_to_ = 0.252 nS/pF).

*I*_sus_:	4-aminopyridine sensitive sustained K^+^ current (*g*_sus_ = 0.02 nS/pF).

*I*_NaK_:	Na^+^/K^+^ pump current (*I*_NaK_= 2.88 pA/pF).

*K*_mKp_ = 1.4 mM: Half-maximal *K*_o_ for *I*_NaK_.

*K*_mNap_ = 14 mM: Half-maximal *Na*_i_ for *I*_NaK_.

*I*_bCa_:	Background Ca^2+^ current (*g*_bCa_ = 0.0006 nS/pF).

*I*_bNa_:	Background Na^+^ current (*g*_bNa_ = 0.00486 nS/pF).

*I*_NCX_:	Na^+^/Ca^2+^ exchanger (NCX) current (*k*_NCX_ = 187.5 pA/pF).

*K*_1ni_ = 395.3: intracellular Na^+^ binding to first site on NCX.

*K*_2ni_ = 2.289: intracellular Na^+^ binding to second site on NCX.

*K*_3ni_ = 26.44: intracellular Na^+^ binding to third site on NCX.

*K*_1no_ = 1628: extracellular Na^+^ binding to first site on NCX.

*K*_2no_ = 561.4: extracellular Na^+^ binding to second site on NCX.

*K*_3no_ = 4.663: extracellular Na^+^ binding to third site on NCX.

*K*_ci_ = 0.0207: intracellular Ca^2+^ binding to NCX transporter.

*K*_co_ = 3.663: extracellular Ca^2+^ binding to NCX transporter.

*K*_cni_ = 26.44: intracellular Na^+^ and Ca^2+^ simultaneous binding to NCX.

*Q*_ci_:= 0.1369: intracellular Ca^2+^ occlusion reaction of NCX.

*Q*_co_=0: extracellular Ca^2+^ occlusion reaction of NCX.

*Q*_n_= 0.4315: Na^+^ occlusion reactions of NCX.

**Ca^2+^ diffusion**

*τ*_difCa_ = 0.04 ms: Time constant of Ca^2+^ diffusion from the submembrane to myoplasm.

*τ*_tr_ = 40 ms: Time constant for Ca^2+^ transfer from the network to junctional SR.

**SR Ca^2+^ ATPase function**

*K*_up_ = 0.6·10^-3^ mM: Half-maximal Ca_i_ for Ca^2+^ uptake in the network SR.

*P*_up,basal_ = 0.012 mM/ms: Rate constant for Ca^2+^ uptake by the Ca^2+^ pump in the network SR.

**RyR function**

*k*_oCa_ = 10 mM^-2^· ms^-1^; *k*_om_ = 0.06 ms^-1^; *k*_iCa_ = 0.5 mM^-1^· ms^-1^ ; *k*_im_ = 0.005 ms^-1^; *EC*_50_SR_ = 0.45 mM; *k*_s_ = 250·10^3^ ms^-1^; *MaxSR =*15; *MinSR =*1; *HSR =* 2.5;

**Ca^2+^ and Mg^2+^ buffering**

*k*_bCM_=0.542 ms^-1^: Ca^2+^ dissociation constant for calmodulin.

*k*_bCQ_=0.445 ms^-1^: Ca^2+^ dissociation constant for calsequestrin.

*k*_bTC_=0.446 ms^-1^: Ca^2+^ dissociation constant for the troponin-Ca^2+^ site.

*k*_bTMC_=0.00751 ms^-1^: Ca^2+^ dissociation constant for the troponin-Mg^2+^ site.

*k*_bTMM_=0.751 ms^-1^: Mg^2+^ dissociation constant for the troponin-Mg^2+^ site.

*k*_fCM_=227.7 mM^-1^· ms^-1^: Ca^2+^ association constant for calmodulin.

*k*_fCQ_=0.534 mM^-1^· ms^-1^: Ca^2+^ association constant for calsequestrin.

*k*_fTC_=88.8 mM/ms: Ca^2+^ association constant for troponin.

*k*_fTMC_=227.7 mM/ms: Ca^2+^ association constant for the troponin-Mg^2+^ site.

*k*_fTMM_=2.277 mM/ms: Mg^2+^ association constant for the troponin-Mg^2+^ site.

*TC*_tot_=0.031 mM: Total concentration of the troponin-Ca^2+^ site.

*TMC*_tot_=0.062 mM: Total concentration of the troponin-Mg^2+^ site.

*CQ*_tot_=10 mM: Total calsequestrin concentration.

*CM*_tot_=0.045 mM: Total calmodulin concentration.

**FORMULATIONS: ELECTROPHYSIOLOGY**

**Membrane potential, *V*_m_ (variable *y_15_*)**

*dV_m_*/*dt = -* (*I*_CaL_ + *I*_CaT_ + *I*_f_ + *I*_st_ + *I*_Kr_ + *I*_Ks_ + *I*_to_ + *I*_sus_ + *I*_NaK_ + *I*_NCX_ + *I*_bCa_ + *I*_bNa_+ *I*_KACh_) /*C*_m_

**Gating variables *(y_16_ - y_29_)* and their differential equations**

*dy_i_/dt = (y_i,∞_ -* *y)*/*τ_yi_*

(*y_i_ =* *d*_L_, *f*_L_, *f*_Ca_, *d*_T_, *f*_T_, *p*_aF_, *p*_aS_, *p*_i_, *n*, *q*, *r*, *y*, *q*_a_, *q*_i_)

*τ_yi_*: Time constant for a gating variable *y_i_*.

*α_yi_* and *β_yi_*: Opening and closing rates for channel gating.

*y_i_,*_∞_: Steady-state curve for a gating variable *y_i_*.

**Ion currents**

**L-type Ca^2+^ current (*I*_CaL_),** based on formulations of Kurata et al. (Kurata et al., 2002) that include Ca^2+^ dependent *I*_CaL_ inactivation.

*I*_CaL_=*C*_m_·*g*_CaL_ ·(*V_m_- E*_CaL_)·*d*_L_·*f*_L_·*f*_Ca_

*d*_L,∞_ =1/{1+ exp[-(*V_m_*- *V*_½,d_ )/ *K*_d_ ]}

*f*_L,∞_ =1/{1+exp[(*V_m_* - *V*_½,f_ )/ *K*_f_ ]}

*α*_dL_ = -0.02839·(*V_m_*+ 35)/ {exp[-(*V_m_*+35)/2.5] - 1} -0.0849 · *V_m_* / [exp(-*V_m_*/4.8)- 1]

*β*_dL_ = 0.01143 · (*V_m_* - 5)/ {exp[(*V_m_*- 5)/2.5] -1}

*τ*_dL_ =1/(*α*_dL_ + *β*_dL_)

*τ*_fL_ = 257.1 · exp{-[(*V_m_*+ 32.5)/13.9]^2^ }+ 44.3

*f*_Ca,∞_ =*K*_mfCa_ / (*K*_mfCa_ + *Ca*_sub_)

*τ*_fCa_ = *f*_Ca,∞_ / *α*_fCa_

**T-type Ca^2+^ current (*I*_CaT_),** based on formulations suggested by Demir et al., (Demir et al., 1994) and modified by Kurata et al. (Kurata et al., 2002).

*I*_CaT_ = *C*_m_∙*g*_CaT_ ∙(*V_m_*- *E*_CaT_)∙*d*_T_∙*f*_T_

*d*_T,∞_ =1/ {1 + exp[-(*V_m_* + 26.3)/6.0]}

*f*_T,∞_ = 1/{1+ exp[(*V_m_*+ 61.7)/5.6]}

*τ*_dT_ = 1/{1.068∙exp[(*V_m_*+ 26.3)/30] + 1.068∙exp[-(*V_m_* + 26.3)/30]}

*τ*_fT_ =1/{0.0153∙exp[- (*V_m_* + 61.7)/83.3] + 0.015∙exp[(*V_m_*+ 61.7)/15.38]}

**Rapidly activating delayed rectifier K^+^ current (*I*_Kr_)**, based on formulations suggested by Zhang et al. (Zhang et al., 2000) and modified by Kurata et al. (Kurata et al., 2002).

*I*_Kr_ = *C*_m_∙*g*_Kr_ ∙(*V_m_* - *E*_K_)∙(0.6∙ *p*_aF_ + 0.4∙ *p*_aS_) ∙ *p*_i_

*p*_a,∞_ =1/ {1 + exp[-(*V_m_*+23.2)/10.6]}

*p*_i,∞_ = 1/ {1 + exp[(*V_m_* + 28.6)/17.1]}

*τ*_paF_ = 0.84655354/[0.0372 ∙ exp(*V_m_*/15.9) + 0.00096 ∙ exp(-*V_m_* /22.5)]

*τ*_paS_ = 0.84655354/[0.0042 ∙ exp(*V_m_* /17.0) + 0.00015 ∙ exp(-*V_m_* /21.6)]

*τ*_pi_ = 1/[0.1 ∙ exp(-*V_m_*/54.645) + 0.656 ∙ exp(*V_m_*/106.157)]

**Slowly activating delayed rectifier K^+^ current (*I*_Ks_)**, based on formulations suggested by Zhang et al. (Zhang et al., 2000).

*I*_Ks_ = *C*_m_∙*g*_Ks_ ∙(*V_m_* - *E*_Ks_)∙ *n*^2^

*α*_n_ = 0.014/ {1 + exp[-(*V_m_*- 40)/9]}

*β*_n_ = 0.001 ∙ exp(-*V_m_*/45)

*n*_∞_ = *α*_n_/(*α*_n_ + *β*_n_)

*τ*_n_ =1/(*α*_n_ + *β*_n_)

**4-aminopyridine-sensitive currents (*I*_4AP_ =*I*_to_ *+ I*_sus_),** based on formulations suggested by Zhang et al. (Zhang et al., 2000).

*I*_to_ = *C*_m_ ∙ *g*_to_ ∙ (*V_m_*- *E*_K_) ∙ *q*∙ *r*

*I*_sus_ = *C*_m_ ∙ *g*_sus_ ∙ (*V_m_* - *E*_K_) ∙ *r*

*q*_∞_ =1/{1 + exp[(*V_m_*+ 49)/13]}

*r*_∞_ =1/{1 + exp[-(*V_m_* - 19.3)/15]}

*τ*_q_ = 39.102/{0.57∙exp[-0.08∙(*V_m_*+44)]+0.065∙exp[0.1∙(*V_m_*+45.93)]}+ 6.06

*τ*_r_ =14.40516/{1.037∙exp[0.09∙(*V_m_*+30.61)]+0.369∙exp[-0.12∙(*V_m_*+23.84)]}+ 2.75352

**Hyperpolarization-activated, “funny” current (*I*_f_)**, based on formulations of Wilders at al. (Wilders et al., 1991) and Kurata et al.(Kurata et al., 2002).

*I*_f_ = *I*_fNa_+ *I*_fK_

*y*_∞_ = 1/{1 + exp[(*V_m_* - *V*_If,1/2_) /13.5]}

τ_y_ = 0.7166529/{exp[-(*V_m_*+ 386.9)/45.302] + exp[(*V_m_* - 73.08)/19.231]}

*I*_fNa_ = *C*_m_∙0.3833 ∙*g*_If_ ∙(*V_m_* - *E*_Na_)∙*y*^2^

*I*_fK_ = *C*_m_∙0.6167 ∙ *g*_If_ ∙(*V_m_* - *E*_K_)∙*y*^2^

**Sustained inward current (*I*_st_),** based on formulations of Shinigawa et al. (Shinagawa et al., 2000) which were adopted for rabbit SANC by Kurata et al. (Kurata et al., 2002).

*I*_st_ = *C*_m_ ∙*g*_st_ ∙ (*V_m_* - *E*_st_)∙ *q*_a_∙ *q*_i_

*q*_a,∞_ =1/{1 + exp[-(*V_m_* + 57)/5]}

*α*_qa_ =1/{0.15 ∙ exp(-*V_m_*/11) + 0.2 ∙ exp(-*V_m_*/700)}

*β*_qa_ =1/{16 ∙ exp(*V_m_*/8) + 15 ∙ exp(*V_m_*/50)}

τ_qa_ =1/(*α*_qa_ + *β*_qa_)

*α*_qi_ =1/{3100 ∙ exp(*V_m_*/13) + 700 ∙exp(*V_m_*/70)}

*β*_qi_ =1/{95∙ exp(-*V_m_*/10) + 50 ∙exp(-*V_m_*/700)} + 0.000229/[1 + exp(-*V_m_*/5)]

*τ*_qi_ =6.65/(*α*_qi_ + *β*_qi_)

*q*_i,∞_ =*α*_qi_ /(*α*_qi_ + *β*_qi_)

**Na^+^-dependent background current (*I*_bNa_)**

*I*_b,Na_ =*C*_m_∙*g*_bNa_∙(*V_m_* - *E*_Na_)

**Na^+^-K^+^ pump current (*I*_NaK_),** based on formulations on Kurata et al. (Kurata et al., 2002), which were in turn based on the experimental work of Sakai et al. (Sakai et al., 1996) for rabbit SANC.

*I*_NaK_ = *C*_m_ ∙*I*_NaK_ ∙{1+(K_mKp_/K_o_)^1.2^}^-1^ ∙ {1+(K_mNap_/Na_i_)^1.3^} ^-1^ ∙ {1+exp[-(*V_m_*- *E*_Na_+120)/30]}^-1^

**Ca^2+^- background current (*I*_bCa_)**

*I*_bCa_ = *C*_m_∙ *g*_bCa_ ∙(*V_m_* - *E*_CaL_)

**Na^+^-Ca^2+^ exchanger current (*I*_NCX_)**, based on original formulations from Dokos et al. (Dokos et al., 1996).

*I*_NCX_ = *C*_m_ ∙*k*_NCX_ ∙(*k*_21_ ∙ *x*_2_ - *k*_12_ ∙ *x*_1_) / (*x*_1_ + *x*_2_ + *x*_3_ + *x*_4_)

*d*_o_ =1+(*Ca*_o_/*K*_co_)∙{1+exp(*Q*_co_∙*V_m_*/*E*_T_)}+(*Na*_o_/*K*_1no_)∙{1+(*Na*_o_/*K*_2no_)∙(1+*Na*_o_/*K*_3no_)}

*k*_43_ = *Na*_i_/(*K*_3ni_ +*Na*_i_)

*k*_41_ = exp[-*Q*_n_∙*V_m_*/(2*E*_T_)]

*k*_34_ = *Na*_o_/(*K*_3no_+*Na*_o_)

*k*_21_ = (*Ca*_o_/*K*_co_)∙exp(*Q*_co_∙*V_m_*/*E*_T_) /*d*_o_

*k*_23_ = (*Na*_o_/*K*_1no_)∙(*Na*_o_/*K*_2no_)∙(1+*Na*_o_/*K*_3no_)∙ exp[-*Q*_n_∙*V_m_*/(2*E*_T_)]/*d*_o_

*k*_32_ = exp[*Q*_n_∙*V_m_*/(2*E*_T_)]

*x*_1_ = *k*_34_ ∙ *k*_41_ ∙(*k*_23_ + *k*_21_) + *k*_21_ ∙ *k*_32_ ∙(*k*_43_ + *k*_41_)

*d*_i_ = 1+(*Ca*_sub_/*K*_ci_)∙{1+exp(-*Q*_ci_∙*V_m_*/*E*_T_)+*Na*_i_/*K*_cni_}+(*Na*_i_/*K*_1ni_)∙{1+(*Na*_i_/*K*_2ni_)∙(1+*Na*_i_/*K*_3ni_)}

*k*_12_ =(*Ca*_sub_/*K*_ci_)∙exp(-*Q*_ci_∙*V_m_*/*E*_T_)/*d*_i_

*k*_14_ = (*Na*_i_/*K*_1ni_)∙(*Na*_i_/*K*_2ni_)∙(1 +*Na*_i_/*K*_3ni_)∙ exp[*Q*_n_∙*V_m_*/(2*E*_T_)]/*d*_i_

*x*_2_ = *k*_43_∙ *k*_32_ ∙(*k*_14_ + *k*_12_) + *k*_41_∙ *k*_12_ ∙*k*_34_ + *k*_32_)

*x*_3_ = *k*_43_ ∙*k*_14_ ∙(*k*_23_ + *k*_21_) + *k*_12_ ∙*k*_23_∙(*k*_43_ + *k*_41_)

*x*_4_ = *k*_34_ ∙*k*_23_ ∙(*k*_14_ + *k*_12_) + *k*_21_ ∙*k*_14_∙(*k*_34_+ *k*_32_)

**FORMULATIONS: Ca^2+^ CYCLING**

**Ca^2+^ release flux (*j*_SRCarel_) from SR via RyRs**, based on original formulations of Stern et al. (Stern et al., 1999) and modified by Shannon et al. (Shannon et al., 2004)

*j*_SRCarel_ = *k*_s_∙*O*∙(*Ca*_jSR_ - *Ca*_sub_)

*k*_CaSR_ = *MaxSR*- (*MaxSR* - *MinSR*)/ (1 + (*EC*_50_SR_/*Ca*_jSR_)^HSR^)

*k*_oSRCa_ = *k*_oCa_/*k*_CaSR_

*k*_iSRCa_ = *k*_iCa_∙*k*_CaSR_

*dR/dt* = (*k*_im_∙*RI* - *k*_iSRCa_ ∙*Ca*_sub_∙*R*) - (*k*_oSRCa_∙*Ca*_sub_^2^∙*R* - *k*_om_∙*O*)

*dO/dt* =(*k*_oSRCa_∙*Ca*_sub_^2^ ∙*R* - *k*_om_∙*O*) - (*k*_iSRCa_∙*Ca*_sub_∙*O* - *k*_im_∙*I*)

*dI/dt* = (*k*_iSRCa_∙ *Ca*_sub_∙*O* - *k*_im_∙*I*) - (*k*_om_∙*I* - *k*_oSRCa_∙*Ca*_sub_^2^ ∙*RI*)

*dRI/dt* = (*k*_om_∙*I* - *k*_oSRCa_∙*Ca*_sub_^2^∙*RI*) - (*k*_im_∙*RI* - *k*_iSRCa_∙*Ca*_sub_∙*R*)

**Intracellular Ca^2+^ fluxes**

**Ca^2+^ diffusion flux** **(*j*_Ca_dif_)** from submembrane space to myoplasm:

*j*_Ca_dif_ = (*Ca*_sub_ - *Ca*_i_)/τ_difCa_

**The rate of Ca^2+^ uptake (pumping) (*j*_up_)** by the SR, based on formulations of SR Ca^2+^ pump function suggested by Luo and Rudy (Luo and Rudy, 1994).

*j*_up_ = *P*_up_ /(1 + *K*_up_/*Ca*_i_)

**Ca^2+^ flux between (network and junctional) SR compartments (*j*_tr_)**:

*j*_tr_ = (*Ca*_nSR_ – *Ca*_jSR_)/τ_tr_

**Ca^2+^ buffering**

*df*_TC_*/dt* = *k*_fTC_∙*Ca*_i_∙(1 -*f*_TC_) - *k*_bTC_ ∙ *f*_TC_

*df*_TMC_/*dt* = *k*_fTMC_ ∙*Ca*_i_ ∙(1- *f*_TMC_ - *f*_TMM_) - *k*_bTMC_ ∙ *f*_TMC_

*df*_TMM_/*dt* = *k*_fTMM_ ∙*Mg*_i_ ∙(1-*f*_TMC_ - *f*_TMM_)- *K*_bTMM_ ∙ *f*_TMM_

*df*_CMi_/*dt* =	*k*_fCM_ ∙*Ca*_i_ ∙(1- *f*_CMi_) - *k*_bCM_ ∙ *f*_CMi_

*df*_CMs_/*dt* = *k*_fCM_ ∙*Ca*_sub_∙(1 - *f*_CMs_) - *k*_bCM_ ∙ *f*_CMs_

*df*_CQ_/*dt* =	*k*_fCQ_ ∙*Ca*_jSR_∙(1- *f*_CQ_) - *k*_bCQ_ ∙ *f*_CQ_

**Dynamics of Ca^2+^ concentrations in cell compartments**

*dCa*_i_/*dt* =(*j*_Ca_dif_ ∙*V*_sub_ - *j*_up_∙ *V*_nSR_) /*V*_i_ - (*CM*_tot_∙*df*_CMi_*/dt* + *TC*_tot_∙*df*_TC_/*dt* + *TMC*_tot_∙*df*_TMC_*/dt*)

*dCa*_sub_*/dt* = *j*_SRCarel_ ∙*V*_jSR_/*V*_sub_ -(*I*_CaL_+*I*_CaT_+*I*_bCa_-2∙*I*_NCX_)/(2∙F∙*V*_sub_)-(*j*_Ca_dif_ + *CM*_tot_ ∙*df*_CMs_*/dt*)

*dCa*_jSR_/*dt* = *j*_tr_ - *j*_SRCarel_ - *CQ*_tot_ ∙ *df*_CQ_*/dt*

*dCa*_nSR_*/dt* = *j*_up_ - *j*_tr_ ∙*V*_jSR_/*V*_nSR_

**Initial values**

Online Table S1 summarizes all model variables (*y_1_*–*y_29_*) with their initial values.

**Autonomic modulation.**

Please note that *g*_CaL_ and *P*_up_ are different in different SAN modules.

**βAR stimulation**

The effect of βAR stimulation was modelled as previously described in our previous studies (Maltsev and Lakatta, 2010).

***g*_CaL_:**

*g*_CaL_βAR_ = *g*_CaL_∙1.75

***If*:**

*V*_If,1/2_βAR_ = *V*_If,1/2_ +7.8 = -64+7.8 = -56.2 mV

***I_Kr_*:**

*g*_Kr_βAR_ = *g*_Kr_∙1.5 = 0.08113973*1.5 = 0.121709595 nS/pF

***P*_up_:**

*P*_up_βAR_ = *P*_up_∙2

**CR stimulation**

The effect of ChR stimulation with 0.1 μM of ACh was modelled as previously described in our previous studies (Maltsev and Lakatta, 2010). However, the effect on I_CaL_ was changed to Zaza et al. model (Zaza et al., 1996) as described in details below (modified from (Yang et al., 2021)).

***I*_CaL_:**

*g*_CaL_ACh_ =*C*_m_·*g*_CaL_· (1 - *b*_CaL_)

The fractional block (*b*_CaL_) of *I*_CaL_ by CR stimulation was adopted from Zaza et al. model given in Figure 2 legend in (Zaza et al., 1996):

*b*_CaL_ = [ACh]^0.348/(2921^0.348+[ACh]^0.348), where [ACh] is given in μM

Thus, the fractional block in the presence of 0.1 μM of ACh (simulated in the present study) was relatively small: *b*_CaL_(0.1) = 0.02728, i.e. <3%

***I*_f_:**

The shift *s* (in mV) of the *I*_f_ activation curve by ChR stimulation was adopted from Zhang et al. 2002 (Zhang et al., 2002).

*V*_If,1/2_ACh_ = *V*_If,1/2_ + *s*

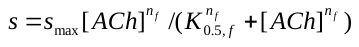

*s*_max_ = -7.2 mV: maximum ACh-induced shift of *I*_f_ half activation voltage.

*n_f_* = 0.69 and *K_0.5,f_* = 12.6 nM: Michaelis-Menten parameters for ACh modulation of *I*_f_.

For 100 nM of ACh: *s_max_* =-5.81 mV and *V*_If,1/2_ACh_ = -64 - 5.81 = - 69.81mV

**Acetylcholine-activated K^+^ current (*I*_KACh_):**

*I*_KACh_ was adopted from Demir et al. 1999 (Demir et al., 1999) (Note *I*_KACh_ = 0 when [ACh]=0)*, g*_KAch_ =0.14241818 nS/pF.

*I*_KACh_ = *a* ⋅ *g*_KACh_ ⋅ (*V*_m_ - *E*_K_)

*beta*=0.001 · 12.32/(1+0.0042/[ACh]) (per ms)

*alfa*= 0.001 · 17⋅exp(0.0133⋅(*V_m_*+40)) (per ms)

*a*_∞_ = *beta* / (*alfa* + *beta*)

*τ*_a_ = 1/(*alfa* + *beta*) (in ms)

The present study did not investigate the kinetics of *I*_KACh_ and the system rtansitions after ACh application. Thus, in our simulations *I*_KACh_ has always reached its steady-state, and *a = a*_∞_. Thus,

*I*_KACh_ = *a*_∞_ ⋅ *g*_KACh_ ⋅ (*V*_m_ - *E*_K_)

***P_up_*:**

ChR stimulation inhibits *P*_up_, with fractional block *b*_up_ formulated as follows:

*P*_up_ACh_ = *P*_up_ · (1 - *b*_up_)

*b_up_ = b_up,max_* ⋅ [ACh]/( *K_0.5,up_* + [ACh])

where *K_0.5,up_* = 90 nM is the [ACh] for half-maximal inhibition and *b_up,max_* = 0.7.

For 100 nM of ACh: *b_up_ =*0.368421052632.
