## Supplementary Table S1 for "A novel conceptual model of heart rate autonomic modulation based on a small-world modular structure and heterogeneous innervation of the sinoatrial node"

**Supplementary Table S1.** Model variables: description and initial values. Our SAN cell model is described as a system of 29 first order differential equations (variables *y_1_ - y_29_*).

| # | Variable | Description | Initial value |
| --- | --- | --- | --- |
| *Ca^2+^ cycling* | | | |
| *y_1_* | *Ca_i_* | [Ca^2+^] in myoplasm, mM | 0.0001 |
| *y_2_* | *Ca_sub_* | [Ca^2+^] in submembrane space, mM | 0.000223 |
| *y_3_* | *Ca*_jSR_ | [Ca^2+^] in the junctional SR (jSR), mM | 0.029 |
| *y_4_* | *Ca*_nSR_ | [Ca^2+^] in the network SR (nSR), mM | 1.35 |
| *y_5_* | *f*_TC_ | Fractional occupancy of the troponin-Ca^2+^ site by Ca^2+^ in myoplasm | 0.02 |
| *y_6_* | *f*_TMC_ | Fractional occupancy of the troponin-Mg^2+^ site by Ca^2^ in myoplasm | 0.22 |
| *y_7_* | *f*_TMM_ | Fractional occupancy of the troponin-Mg^2+^ site by Mg^2+^ in myoplasm | 0.69 |
| *y_8_* | *f*_CMi_ | Fractional occupancy of calmodulin by Ca^2+^ in myoplasm | 0.042 |
| *y_9_* | *f*_CMs_ | Fractional occupancy of calmodulin by Ca^2+^ in submembrane space | 0.089 |
| *y_10_* | *f*_CQ_ | Fractional occupancy of calsequestrin by Ca^2+^ in junctional SR | 0.032 |
| *y_11_* | *R* | RyR reactivated (closed) state | 0.7499955 |
| *y_12_* | *O* | RyR open state | 3.4·10^-6^ |
| *y_13_* | *I* | RyR inactivated state | 1.1·10^-6^ |
| *y_14_* | *RI* | RyR RI state | 0.25 |
| *Electrophysiology* | | | |
| *y_15_* | *V_m_* | Membrane potential, mV | -65 |
| *y_16_* | *d*_L_ | *I*_CaL_ activation | 0 |
| *y_17_* | *f*_L_ | *I*_CaL_ voltage-dependent inactivation | 1 |
| *y_18_* | *f*_Ca_ | *I*_CaL_ Ca^2+^ dependent inactivation | 1 |
| *y_19_* | *p*_aF_ | *I*_Kr_ fast activation | 0 |
| *y_20_* | *p*_aS_ | *I*_Kr_ slow activation | 0 |
| *y_21_* | *p*_i_ | *I*_Kr_ inactivation | 1 |
| *y_22_* | *n* | *I*_Ks_ activation | 0 |
| *y_23_* | *y* | *I*_f_ activation | 1 |
| *y_24_* | *d*_T_ | *I*_CaT_ activation | 0 |
| *y_25_* | *f*_T_ | *I*_CaT_ inactivation | 1 |
| *y_26_* | *q* | *I*_to_ inactivation | 1 |
| *y_27_* | *r* | *I*_to_ and *I*_sus_ activation | 0 |
| *y_28_* | *q_a_* | *I*_st_ activation | 0 |
| *y_29_* | *q_i_* | *I*_st_ inactivation | 1 |
