## Supplementary Figure S1 for "A novel conceptual model of heart rate autonomic modulation based on a small-world modular structure and heterogeneous innervation of the sinoatrial node"

### Slide 1
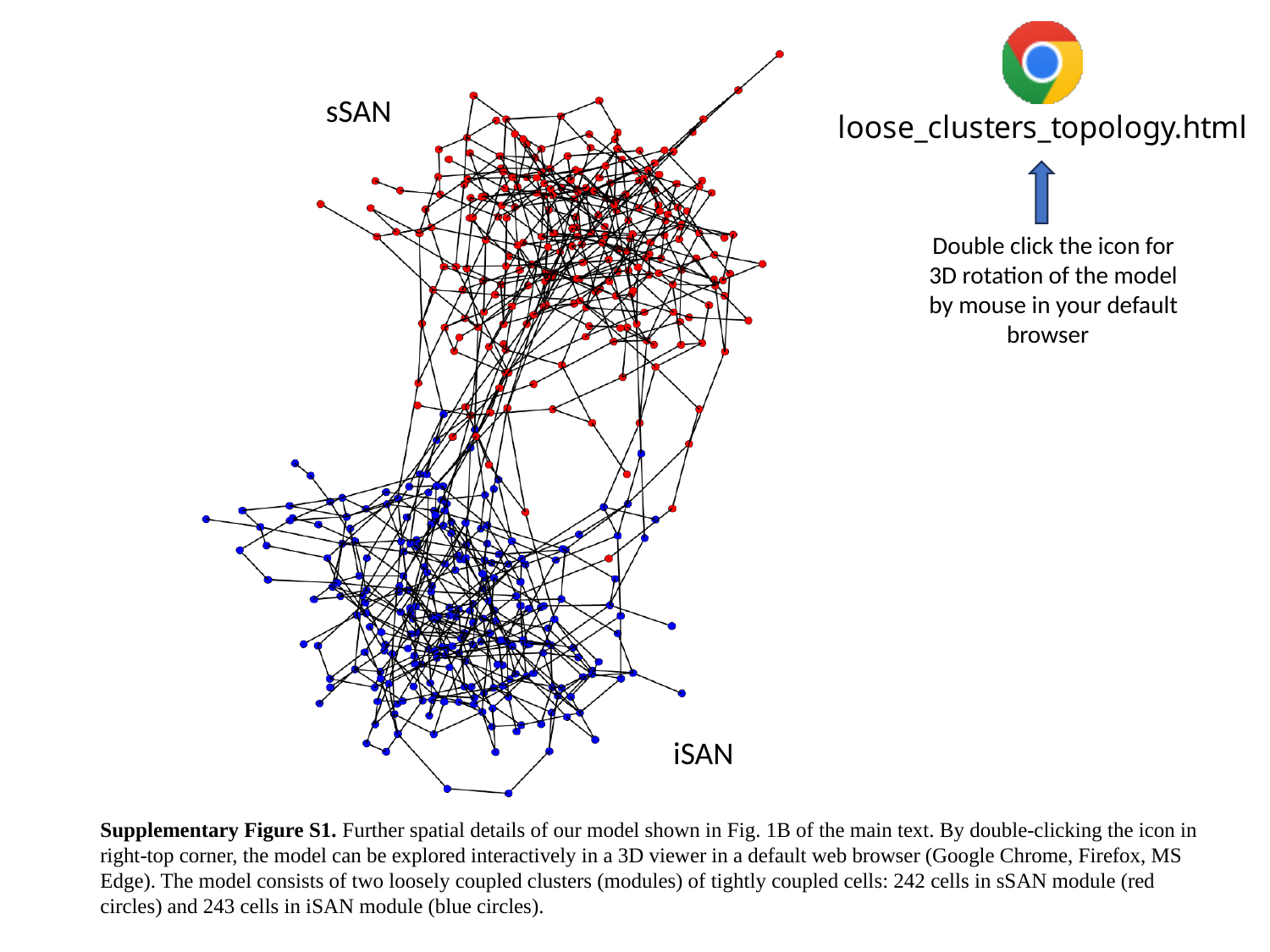

sSAN
Double click the icon for 3D rotation of the model by mouse in your default browser
iSAN
Supplementary Figure S1. Further spatial details of our model shown in Fig. 1B of the main text. By double-clicking the icon in right-top corner, the model can be explored interactively in a 3D viewer in a default web browser (Google Chrome, Firefox, MS Edge). The model consists of two loosely coupled clusters (modules) of tightly coupled cells: 242 cells in sSAN module (red circles) and 243 cells in iSAN module (blue circles).
